# Production of a resilient biosurfactant using agro–wastes and *Paenibacillus dendritiformis* isolated from petroleum-contaminated soil: a break-even analysis at pilot scale

**DOI:** 10.64898/2026.10.19.760049

**Authors:** Gayatri Jeph, Pankaj Goswami, Vanlal Duhsaki, Krupa Chauhan, Ankit Baheti, Mayur Sahani, Shruti Mathur

## Abstract

Wastes from mustard oil mills and dairy were used as partial replacement of pure reagents in modified Bushnell and Haas broth media for production of biosurfactant by *Paenibacillus dendritiformis* isolated from petroleum-contaminated soil. The process was optimised by construction of a response model using Taguchi experimental design, leading to a 28.23% increase in performance characteristics and a 13.23 % increase in yield of biosurfactant. The optimized production medium consisted of carbon @7.9 %, nitrogen @ 1.1% and agitation rate 180 r.p.m. This was provided by adding glucose @ 2%, ammonium nitrate @0.2%, mustard oil seed meal @10% and whey@ 10% in modified Bushnell Haas broth. The biosurfactant produced retained its properties during exposure to a wide range of pH values (5–11), high temperatures (up to 121C), high salinities of up to 5% and also five -fold concentration.

Approximately three-fold increase in yield of the biosurfactant (25g/L to 68.5g/L) was obtained upon scaling-up the production from 100ml to 1500 ml in a lab scale fermenter with agitation @ 150r.p.m. and D.O. levels maintained at 6mg/L.

Power calculations and the cost thereof was calculated (Rs 10.24 /L) and summed up with the cost of raw materials Rs 8.40/L and found to be Rs 18.64/L of product .Since 68.5g product is present in 1L, the cost is calculated to be Rs 271/- per Kg.

Comparison with selling price of currently used chemical surfactants like Linear Alkyl Benzene sulphonic acid (LAB) and sodium Lauryl sulphate and Sodium Lauryl ether sulphate (SLS and SLES) reveals that prices of these products varies from Rs 120 to 600 per kg depending on manufacturer and quality, order quantity.

From the above studies it can be seen that production and use of this multiutility biosurfactant has, besides a clear advantage over chemically produced surfactants in terms of sustainability, a fairly good comparison with selling price of currently used chemical surfactants.

The present study on the production costs (excluding capital costs) successfully demonstrates a break even analysis into the production of biosurfactants.

**Graphical abstract:** 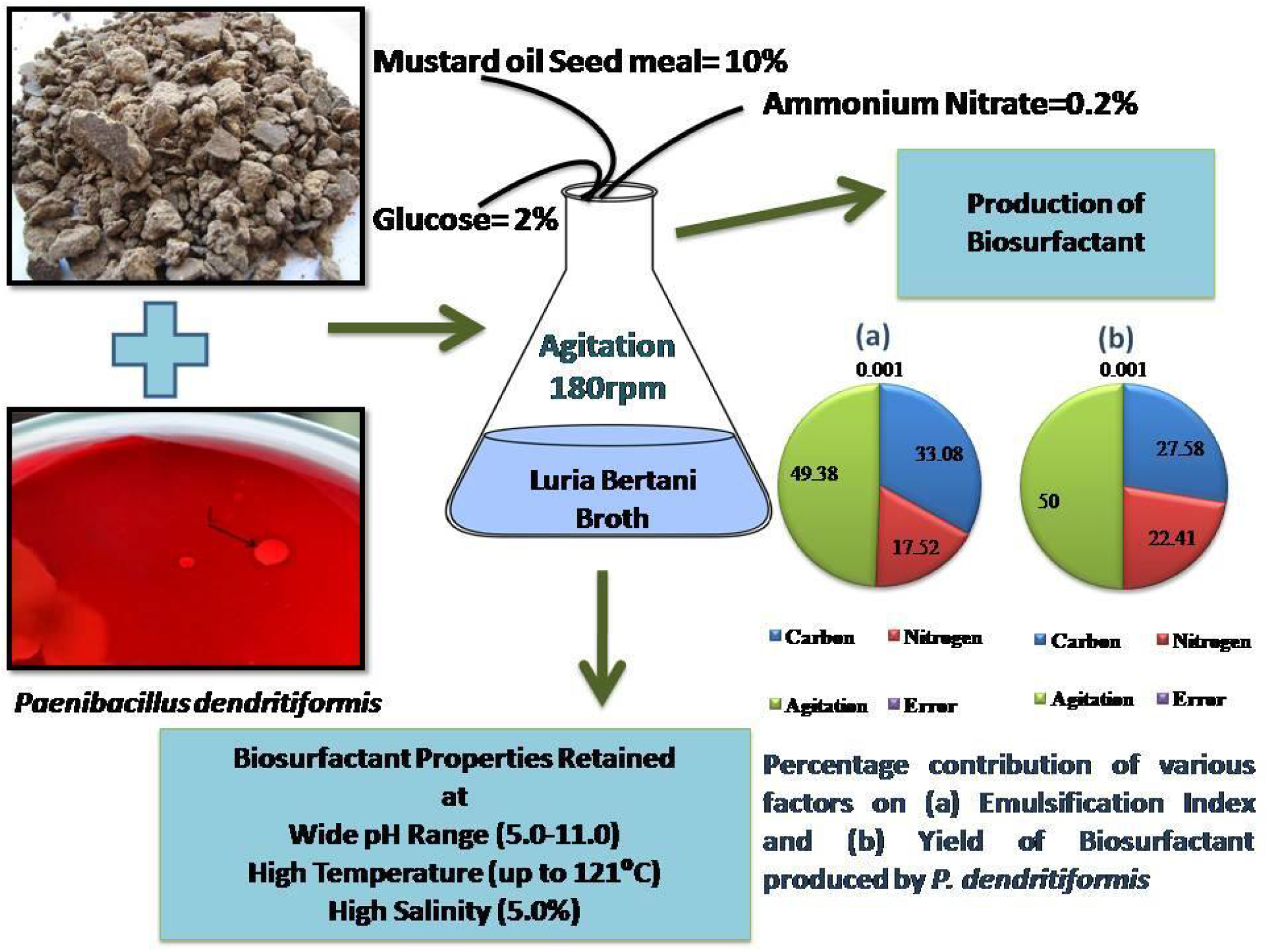

**Statement of Novelty:**

1. Use of mustard oil seed meal as a media supplement for biosurfactant production with enhanced yield and performance characteristics after media optimization has been reported for the first time.
2. The work is novel and pertains to value addition to waste of oil seed industry.
3. The process developed is scalable, commercially viable and both, the process and the product have an inherently good sustainable profile.

## Introduction

Emulsifiers and surfactants are widely used in the pharmaceutical sector, cosmetics industry, soap and detergent industry, textile industry, oilfield chemicals, agrochemicals, food industry, paints & coatings amongst others. About 54% of the total surfactant output is utilized in household/laundry detergents, with only 32% for industrial use. There is a high demand from these end sectors influencing market growth. According to market research sources, the soap/detergent sector contributes maximally to the global surfactants market, due to the increase in awareness of personal and community hygiene. Additionally, the Global Surfactants Market is anticipated to grow at a value CAGR of 5.7%, reaching a valuation of US$ 96.3 billion between the years of 2023 to 2031 [1]. The elastomers and plastics application segment and the oilfield chemicals surfactants segment in the North American region is expected to lose its dominance to Asia-Pacific, owing to the increased off-shore exploration activities and expected commercial production of shale gas by China, which has the largest shale gas reserves [2].

However, UN Environment Assembly in 2016 sounded the alarm bells for policymakers and other stakeholders to understand the critical role of the sound management of chemicals and waste in sustainable development [3].

The global biosurfactants market is projected to grow from $4.18 billion in 2022 to $6.04 billion by 2029, at a CAGR of 5.4% in forecast period, 2022-2029 [4]. The global pandemic harmed communal health and significantly hampered people and several nations worldwide.Since the beginning of the pandemic in December 2019, consumption of a variety of items particularly soaps, disinfectants and hand-sanitizers, spiked.

Biosurfactants (BS) are environmentally friendly options produced by microbial fermentation processes. Surfactant properties enable emulsifying, foaming, detergency and dispersing properties [5]. The fact that only small amounts of BS are needed to reduce surface tension coupled with their little or no toxicity and higher biodegradability makes them environmentally friendly comparing with chemical surfactants. Additionally, their stability at extreme environmental conditions and variety in structural and chemical composition makes them versatile in terms of applicability [6]. The potential of these biosurfactants to be nontoxic and effective in cleaning has been known, leading to a significant increase in demand from cleaning product manufacturers.

Furthermore, it has also been found that biosurfactants are particularly efficient in the green synthesis of nanoparticles, and that they could be utilised for dye decolorization of a widely used textile dye, malachite green, without the use of any adjuvant, catalyst, or alteration in ambient conditions [7]. Chromium is a known contaminant that is produced by a variety of industrial processes such as dyeing, steel manufacture, electroplating, painting, and leather tanning. Several studies have been conducted in order to develop a more efficient and cost-effective technique of chromium removal, and some have indicated that extracted EPS could be used for heavy metal remediation [8].

Owing to the growing awareness towards the use of renewable-based products interest has grown towards the use of BS as environmental friendly alternatives to chemically synthesized surfactants [9]. BS can be obtained either by chemical synthesis from renewable resources, by microbial fermentation processes or by enzymatic syntheses [10]. However biosurfactant have not yet been employed much for use in the industry due to technical and economic inadequacies.

To reduce the cost and increase the yield the right microorganism and media incorporating agro-industrial residues have to be developed because its application provides an alternative way to reduce the production cost and help to solve many environmental hazards [11]. The main problem in using the alternative substrates as culture medium is to find a waste with the right balance of nutrients that permits cell growth and product accumulation [12].

The world production of oils and fats is about 2.5-3 million tons, 75% of which are derived from plants and oil seeds. India has a high capacity to generate vegetable oil and there are significant waste from industries associated with soybean, sunflower, olive, groundnut, rapeseed, safflower, sesame, coconut, palm and mustard oils refining. Mustard oil is the third largest source of oil in India. The extraction of oil from mustard seeds results in generation of large amounts of wastes and their disposal is a serious problem. Mustard seed meal is the solvent extracted oil seed cake. However, certain plants within the Brassicaceae family cannot be used as additive in animal feeds or fertiliser because of their biocidal properties.When hydrolyzed, their glucosinolates are converted into biocidal chemicals such as isothiocyanates, nitriles, and ionic thiocyanates. They suppress a number of soil pathogens, insects, and weeds and are therefore used as biofumigants and biopesticides [13,14,16, 17, 18,19].

To add to this, the fact that, the cultivation of additional oilseed varieties, at the large scale necessary to meet worldwide demands including that to supplement fossil fuels, may saturate existing markets for these oilseed meal co-products . These would therefore be available as cheap raw materials for media supplementation and would additionally offer a promising strategy for development of treatment methods prior to disposal of this solid waste [19].

Biosurfactant production is known to be strongly influenced by medium components such as carbon sources, nitrogen sources, growth factors, and inorganic salts and environmental factors and growth conditions such as pH, temperature, agitation and oxygen availability also affect biosurfactant production through their effects on cellular growth or activity byproducts for biosurfactant production. Traditional methods of optimization involve changing one variable while fixing the others at a certain level. This single dimensional search is laborious, time consuming and incapable of reaching a true optimum due to interactions among variables [20].

A statistical optimization strategy through factorial design developed for the optimization of process explores the relationships between several exploratory variables and one or more response variables.[21-23]

The described C and N sources in most studies such as glucose, glycerol, acetates and other organic acids are quite expensive and cannot reduce the cost of biosurfactant production. An approach to lessen the cost is provided by partial or complete replacement of pure reagents with agricultural mixtures .Few attempts at using wastes for biosurfactant production for example, peat hydrolysate, olive oil mill effluent, lactic whey, soybean, potato process effluent and molasses [24-28]. No studies have so far been made using mustard oil seed meal.

The present work was undertaken to study the use of mustard oil seed meal as a possible additive to medium for production of biosurfactants. The study has approached the problem of media optimization for the production of biosurfactant utilizing, both, the conventional as well as the statistical approach making use of waste product of mustard oil mills. Further, the study has projected the lab-scale findings to pilot scale with break-even analysis as a step towards commercialization.

## Materials and Methods

### 2.1 Collection of Waste Sample

Oil seed meal from a mustard oil mill was used as agro-waste (A.W.). Samples were collected from a small-scale mustard oil mill in Neem-Ka-Thana, a village in the vicinity of Jaipur city in Rajasthan, India. Whey was obtained from the local dairy.

### 2.2 Organism and Growth Conditions

A bacterium capable of growth on the waste was isolated from oil contaminated site from the soil of a motor garage in Jaipur, Rajasthan, India. It was identified by 16S ribosomal RNA gene sequencing as *Paenibacillus dendritiformis*.[19]. Isolation was performed using dilution method on Luria Bertani agar. Optimal pH and temperature for growth was determined by growth of bacterial culture in media with pH varying between 3.0 and 10.0 and temperature varying between 4ºC to 60ºC. It was subsequently grown on broth supplemented with varying concentrations of waste and at optimal pH and temperature .

For the production of EPS, the bacterial strains were grown in modified Bushnell and Haas Broth (BHB) for 5 days at 37 °C in shaking condition at 150 r.p.m [29]. Growth was followed by measuring absorbance at 540 nm of supernatant after centrifugation of culture at 4000 r.p.m.to remove the suspended waste material.

### 2.3 Optimization of Production Medium

Initially the media optimization was conducted by conventional optimization, in a series of experiments changing one variable at a time. The seven factors that were chosen to determine optimum conditions for growth and biosurfactant production are as given: Carbon sources (C), Nitrogen sources (N),Agro-waste (A.W.) concentration were (0%, 1%,2%,5%, 10% and 20%), pH(3.0,5.0,7.0 and 10.0) and temperature (4°C,18°C,22°C,37°C,40° and 60°C), agitation speed (120,150,180 and 210 r.p.m) and time of incubation (up to 5 days).

The pH was maintained using citrate buffer for pH (3.0 and 5.0) and tris for alkaline pH (8.0 and 11.0).

The carbon sources used were glucose (2%), n-hexadecane (3%), glycerol (3%) and starch (2%) with NH_4_Cl (0.1%) as nitrogen source.

For evaluation of the most appropriate nitrogen source for the production of biosurfactant, NH_4_Cl (0.1%), NaNO_3_ (0.1%) and NH_4_NO_3_ (0.2%) were employed with carbon source glucose (2%).

Conditions that were found to be optimal by conventional method were used to determine the various levels interaction between variables using Taguchi design as follows:

Three different levels of medium components and rate of agitation were selected for optimization as given in Table1.Design of experiments as per Taguchi design is shown in Table 2.

**Table 1.** Levels of medium components selected for optimization.

| S.No. | Components | Level 1 | Level 2 | Level 3 |
|---|---|---|---|---|
| 1 | Carbon (w/v%) | 7.9 | 14.78 | 17 |
| 2 | Nitrogen (w/v%) | 0.16 | 0.52 | 1.1 |
| 3 | Agitation (rpm) | 120 | 180 | 210 |

The percentage C and N content were provided using glucose, glycerol, A.W. and NaNO_3_, NH_4_Cl and A.W., respectively.

**Table 2.** L9 Orthogonal array by Taguchi design.

| Experiment | Nitrogen content (%) | Agitation (rpm) | Carbon content (%) |
|---|---|---|---|
| 1 | 0.52 | 120 | 7.9 |
| 2 | 1.1 | 180 | 14.78 |
| 3 | 0.16 | 210 | 17 |
| 4 | 1.1 | 120 | 14.78 |
| 5 | 0.52 | 180 | 17 |
| 6 | 1.1 | 210 | 7.9 |
| 7 | 0.16 | 120 | 17 |
| 8 | 0.52 | 180 | 7.9 |
| 9 | 1.1 | 210 | 14.78 |
| Whey was added at varying concentrations (A-G) to the above optimized medium containing 10% oil-seed meal as follows: |  |  |  |
| Composition | Whey (ml) | Modified BH broth (ml) |  |
| A | 70 | 30 |  |
| B | 60 | 40 |  |
| C | 50 | 50 |  |
| D | 40 | 60 |  |
| E | 30 | 70 |  |
| F | 20 | 80 |  |
| G | 10 | 90 |  |

Statistical analysis was done using SPSS Version 20, IBM.

All media were sterilized by autoclaving at 15 psi pressure at 121°C for 1h. Cultures were incubated at 37°C, pH 7.0 for 6d. Statistical analysis was done using SPSS Version 20, IBM.

Bacterial growth was estimated by measurement of absorbance at a wave length of 660 nm. EI% was measured in cell free supernatant obtained by centrifugation at 10,000 X g for 20 minutes.

### 2.4 Crude biosurfactant recovery

Cell free broth was obtained by centrifugation at 10,000X g for 10 m.Biosurfactant was recovered from cell free supernatant by precipitation after adjusting pH to 2.0 using 6N HCl and keeping it at 4°C overnight. The precipitate thus obtained was pelleted at 10,000 rpm for 20 minutes, dried and weighed [30]. This was used to calculate yield of BS per gram of wet biomass.

### 2.5 Surface activity measurement

#### 2.5.1 Surface tension reduction measurements

Surface Tensiometer by Du Nuoy method was used to measure surface tension[31]

#### 2.5.2 Emulsification Index (E1%)

The emulsification index was determined by using a modification of the method described by S. Iqbal and his team[32].

### 2.6 Stability of Biosurfactant

Stability of biosurfactant was studied under varying conditions of pH, temperature and salinity by measuring surface tension and EI(%).

#### 2.6.1 Temperature Stability

Cell free supernatant containing crude biosurfactant was subjected to different temperatures (- 4°C, 10°C, 25°C, 37°C, 60°C, 80°C and 121°C) for a period of 1 hour. Emulsification activity was then estimated.

#### 2.6.2 pH Effect

Cell free supernatant containing crude biosurfactant was subjected to different pH (3.0, 5.0, 7.0, 8.0 and 11.0) for a period of 1 hour. Surface activity was then estimated as described in 2.5 above.

#### 2.6.3 Salinity effect

Cell free supernatant containing crude biosurfactant was subjected to different salinity range (3.5% and 5%)) for a period of 1 hour. Surface activity was then estimated as described in 2.5 above.

### 2.7 Test for toxicity

To determine and validate the safety of the compound produced, various aliquot of cell free supernatant containing 3.2% of product were used to study the toxicity of the compound. This was done twice, once with an exposure duration of one hour and again after 48 hours of exposure.

### 2.8 Scale-up

The optimized conditions as obtained by the above experiments were employed in the fermentor in a volume of 1500ml.The fermentor was operated at temperature of 37°C agitation speed of 180 rpm aeration of 1.5 v.v.m. (without inoculation for first 36 hours) to obtain dissolved oxygen at 6.0ppm [33, 34].

Yield was determined as given in 2.4 above.

### 2.8 Input Power Analysis

To calculate the input power, the method by Senda et. Al was used [35], and Reynolds number was determined by calculating the viscosity of liquid [36].

Reynolds number was calculated using the following relationship:

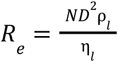

Where η_l = viscosity of liquid N = Rotational Speed

ρ_*l*_ = density of liquid

D = Diameter of stirrer

Viscosity of sample was found by measuring terminal velocity of a given spherical body using the formula below:

viscosity 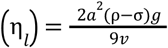

where

Density of steel ball (ρ)=7860 Kg/m^3

Density of sample (σ)=1132 Kg/m^3

Terminal velocity (v)=0.023 m/s

Acceleration due to gravity (g)=10 m/s^2

Radius of spherical steel ball (a)=0.75 cm

Putting all the values in the above formula, the value of viscosity, η= 3.778 Ns/m ^2^ and *R*_*e*_ = 131. 17

Since power curve is reported to be inversely proportional to the Reynolds number at small Reynolds numbers [34].

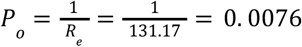

Power input was calculated using the following relationship:

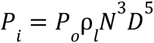

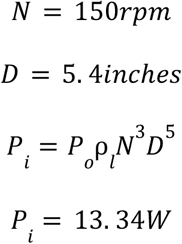

The power consumed for agitation of media at a speed of 150 r.p.m. for a period of 6 days was found to be 1.280 kWh (1.280 Units) per litre. According to the prevailing cost of electricity @ Rs 8 per unit, the cost of operation was calculated to be Rs 10.24 per litre.

## Results and Discussion

The best growth was found to be on media supplemented with glucose (2%) as Carbon source (Fig.1a), ammonium nitrate (0.2%) as nitrogen source(Fig 1b), 210 r.p.m. agitation rate(Fig 1c), pH of 7.0 (Fig 1 d) and temperature of 37°C (Fig 1e). The bacterial strain did not show growth at pH 3.0, 5.0 and 10.0. and at 4°C, 40°C and 60°C. Basal level of growth was however observed at 18°C and 22°C.

**Fig. 1.**
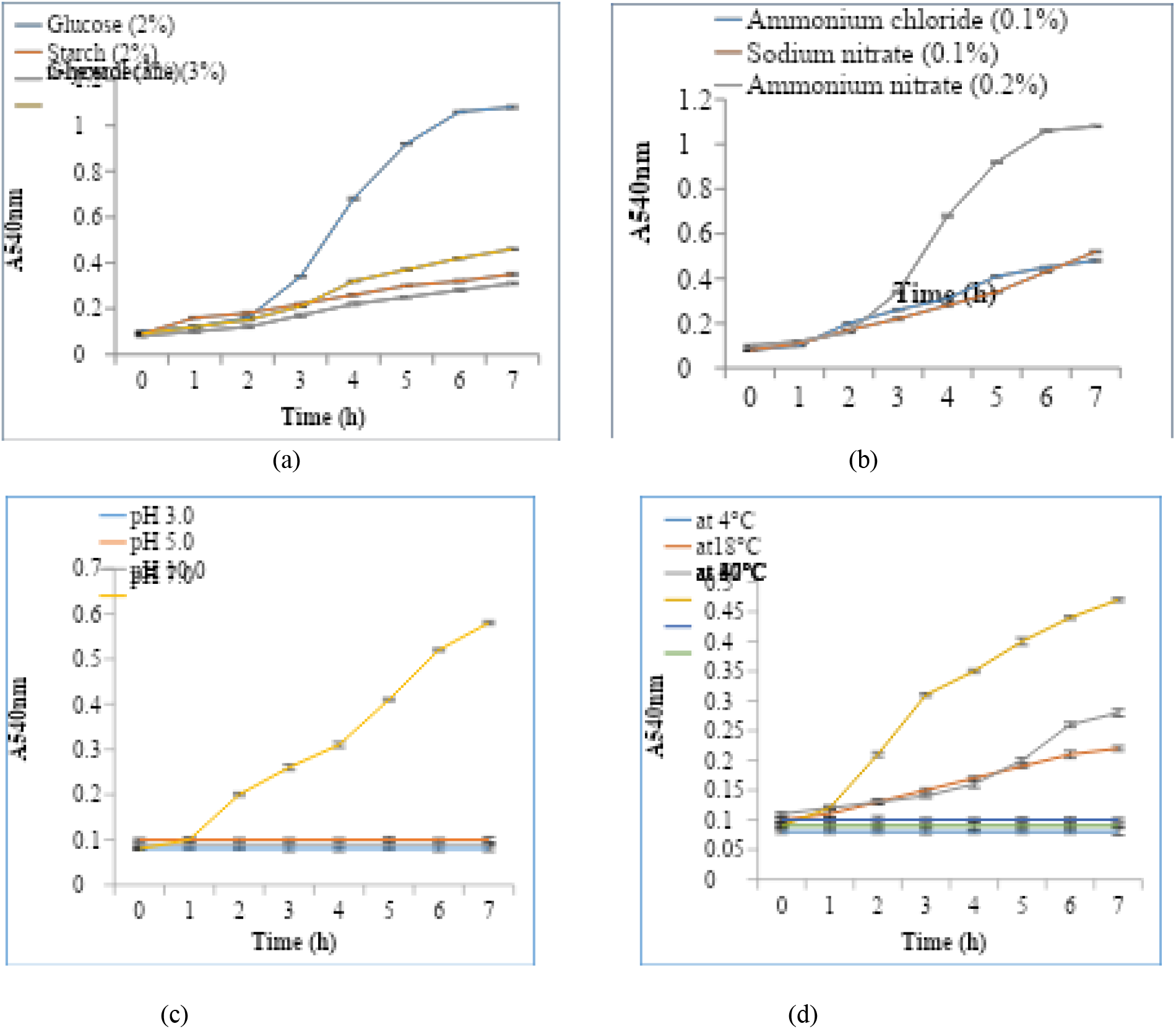
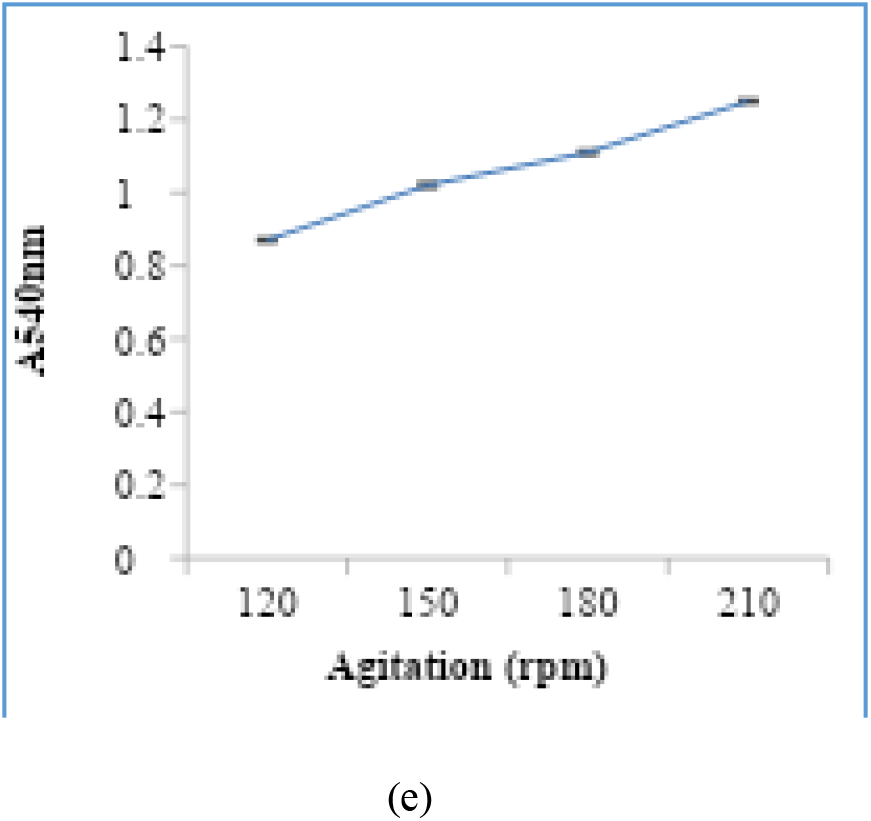
Growth of *Paenibacillus dendritiformis* under various regimes (a) Carbon sources (b) Nitrogen source (c) pH (d) temperature and (e) rates of agitation.

The conditions so optimized were used to amend media with A.W. at varying concentrations and assess BS yield and activity . It was found that growth was best at 10% A.W. amendment (Fig 2). The yield (Table 3) and EI (Fig 3a) also remarkably increased with growth on media containing A.W. Similar increase in EI was observed when compared with various N sources.

**Table 3.**
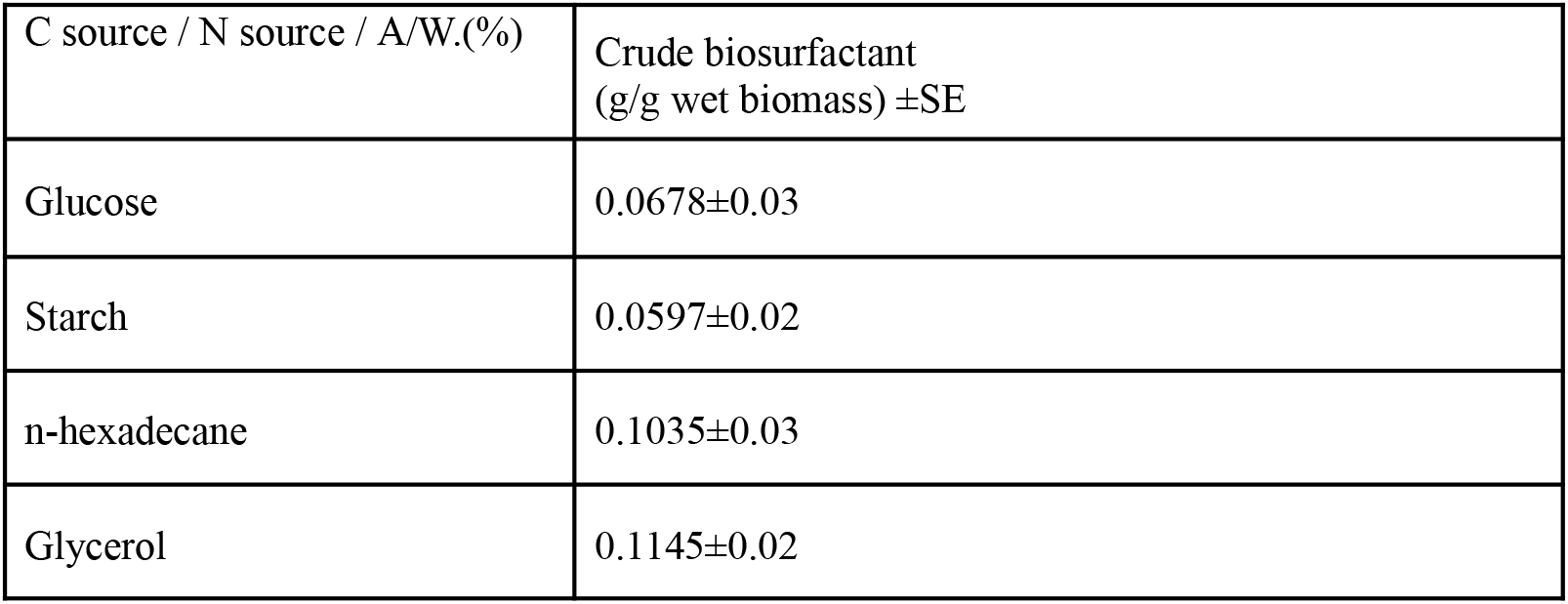
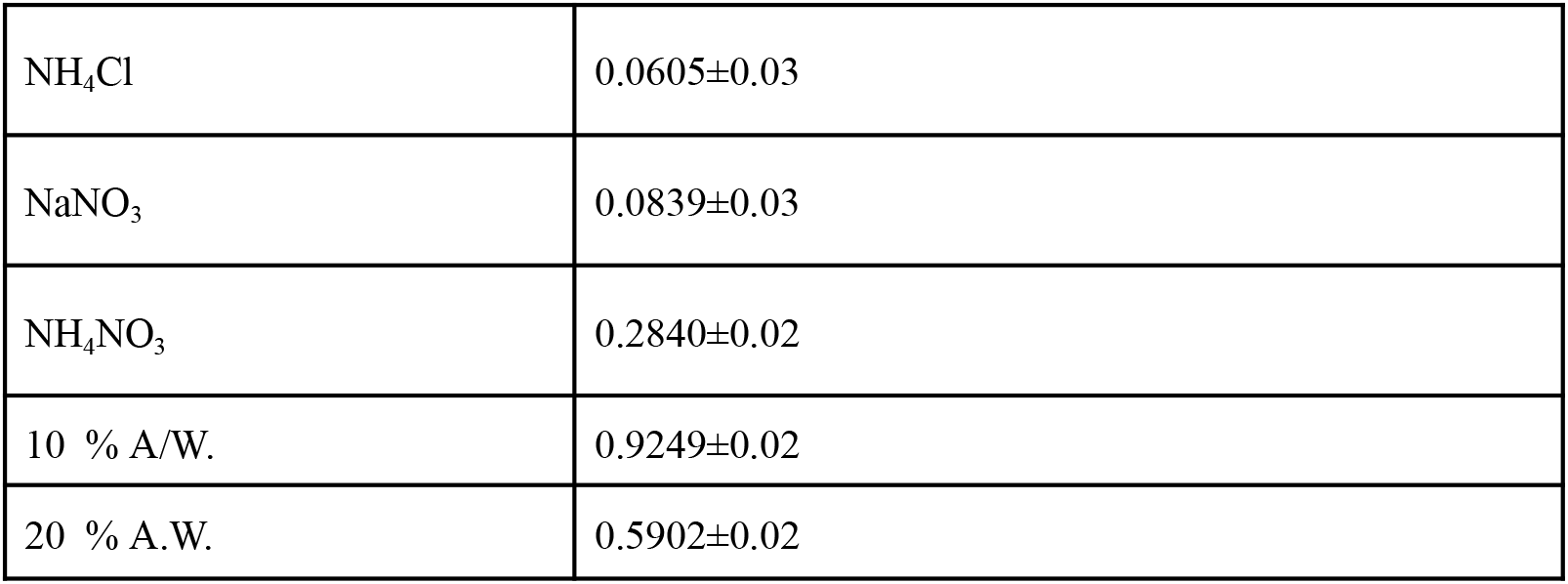
Yield of BS following growth on media with various C, N sources and A.W.

| C source / N source / A/W.(%) | Crude biosurfactant<br>(g/g wet biomass) ±SE |
|---|---|
| Glucose | 0.0678±0.03 |
| Starch | 0.0597±0.02 |
| n-hexadecane | 0.1035±0.03 |
| Glycerol | 0.1145±0.02 |
| NH<sub>4</sub>Cl | 0.0605±0.03 |
| NaNO<sub>3</sub> | 0.0839±0.03 |
| NH<sub>4</sub>NO<sub>3</sub> | 0.2840±0.02 |
| 10 % A/W. | 0.9249±0.02 |
| 20 % A.W. | 0.5902±0.02 |

**Fig. 2.**
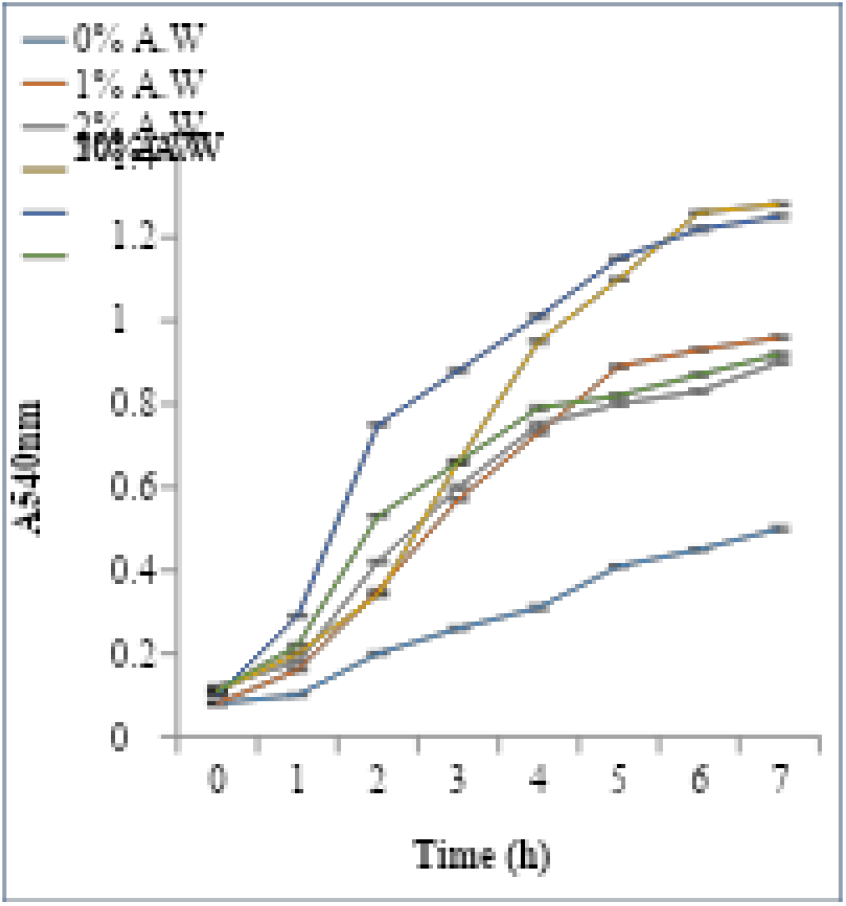
Growth of *P. dendritiformis* at varying concentration of A.W. amended media.

The total C and N contents with media amendments at different levels were then incorporated into growth media at various levels (Table 1) and results of L9 orthogonal Taguchi design were used to obtain the predicted conditions for best BS yield and EI activity (Table 4).

**Table 4.** Various parameters of BS produced by *P.dendritiformis* as per L9 design.

| Group | Carbon<br>Content<br>(%) | Nitrogen<br>Content<br>(%) | Agitation<br>(rpm) | EI<br>(%) | BS yield<br>(g/g wet biomass) |
|---|---|---|---|---|---|
| 1 | 7.9 | 0.52 | 120 | 60 | 0.060 |
| 2 | 14.78 | 1.1 | 180 | 56.88 | 0.34 |
| 3 | 17 | 0.16 | 210 | 65 | 0.19 |
| 4 | 14.78 | 1.1 | 120 | 72.33 | 1.13 |
| 5 | 17 | 0.52 | 180 | 60.5 | 0.831 |
| 6 | 7.9 | 0.16 | 210 | 62 | 0.08 |
| 7 | 17 | 0.16 | 120 | 62.5 | 0.58 |
| 8 | 7.9 | 0.52 | 180 | 70.96 | 1.017 |
| 9 | 14.78 | 1.1 | 210 | 68 | 0.686 |

One way ANOVA showed that nearly 50% contribution for EI and yield was that of rate of agitation. (Fig 4 a and b).

**Fig. 3.**
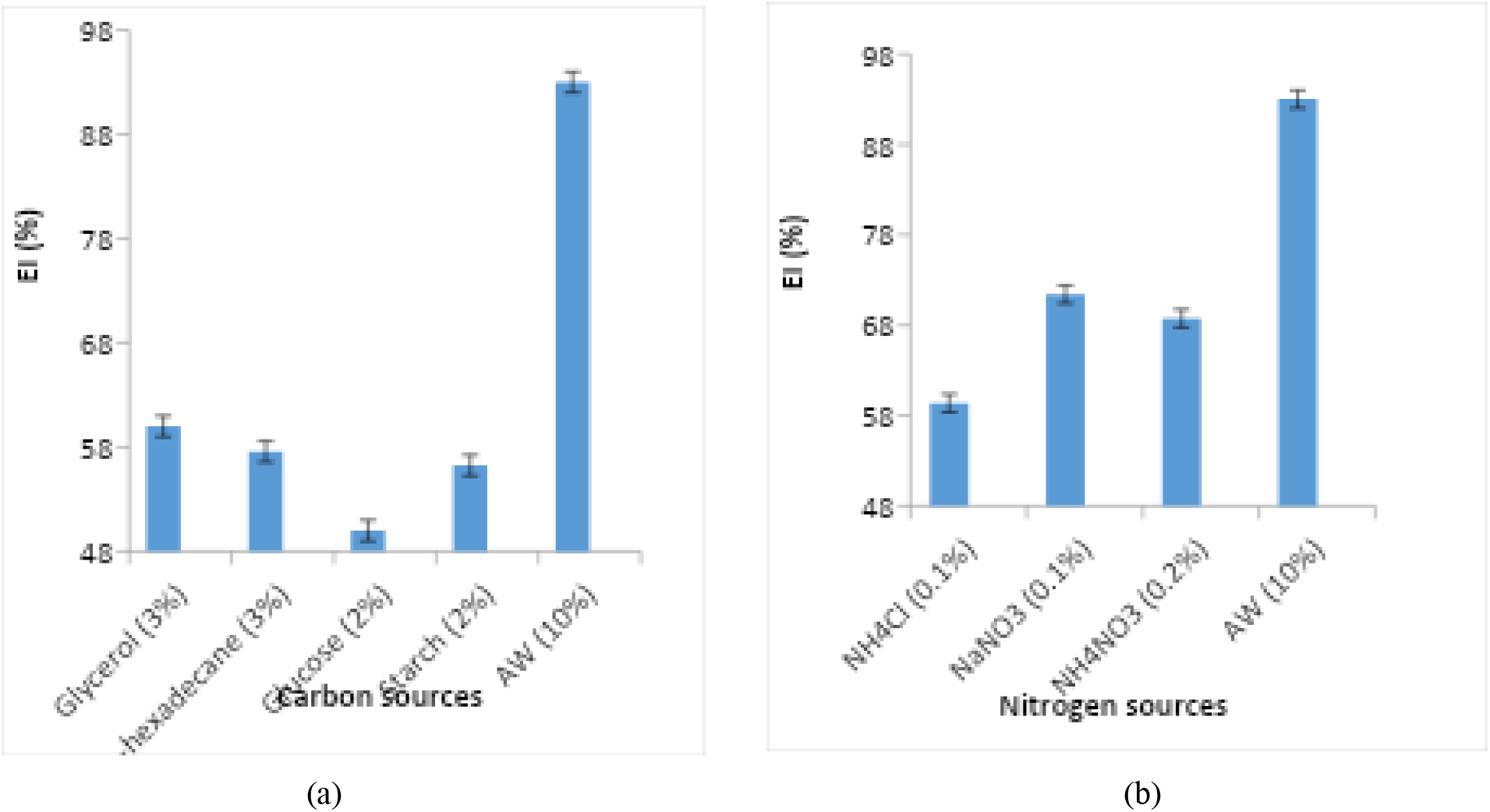
EI of BS obtained following growth on various C sources (a) and N sources (b) *vis a vis* on media amended with A.W.

**Fig. 4.**
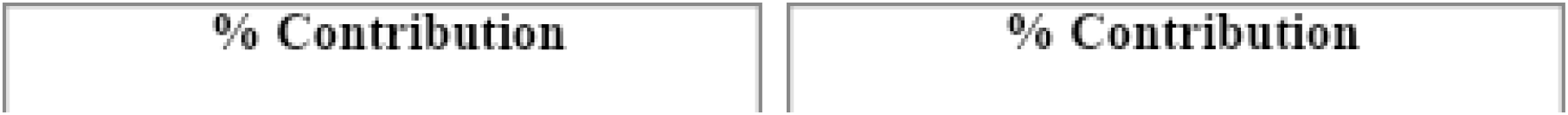
Percentage contribution of various factors on EI (a) and Yield (b) of BS produced by *P. dendritiformis* (Error signifies unexplained variation as per Taguchi experimental design).

Taguchi method of optimization which involves establishment of large number of experimental situations described as orthogonal arrays (OA) to reduce experimental errors and to enhance efficiency and reproducibility of the laboratory experiments shows the combined effect of many factors simultaneously (Y. Abdel-Fattah et al., 2005; S.K. Jha et al., 2014).

Based on minimum SD values (Table 5a) it was statistically predicted that maximum BS yield would be obtained on media of composition : carbon content 7.9 %, nitrogen content 1.1% and agitation rate 180 r.p.m. Also, media composition that would produce BS having best EI activity was as follows: carbon content 7.9%, nitrogen 1.1% and agitation rate 210 r.p.m.(Table 5b). These media conditions are satisfied by amendment of media with A.W.at 10% and whey.

**Table 5.**
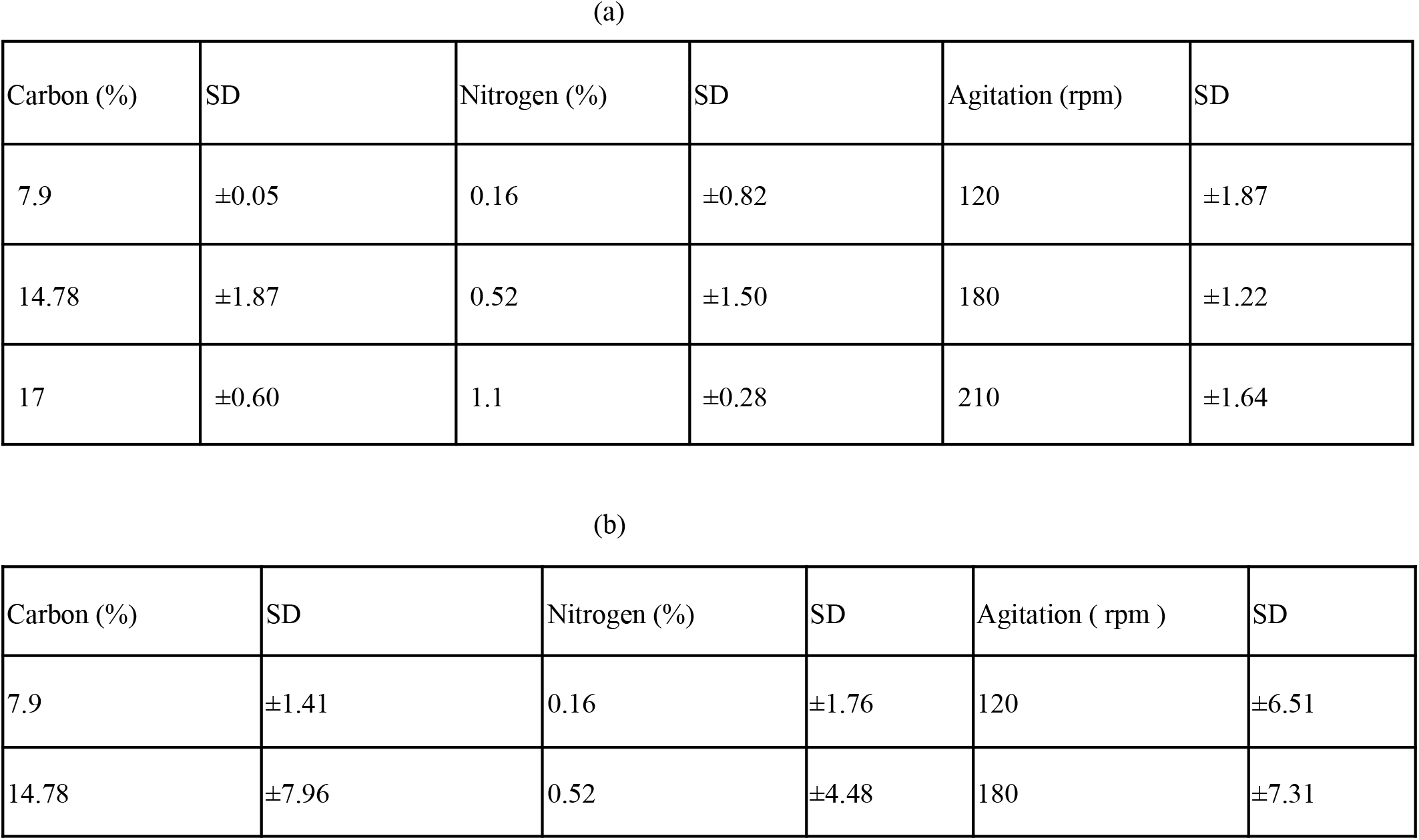
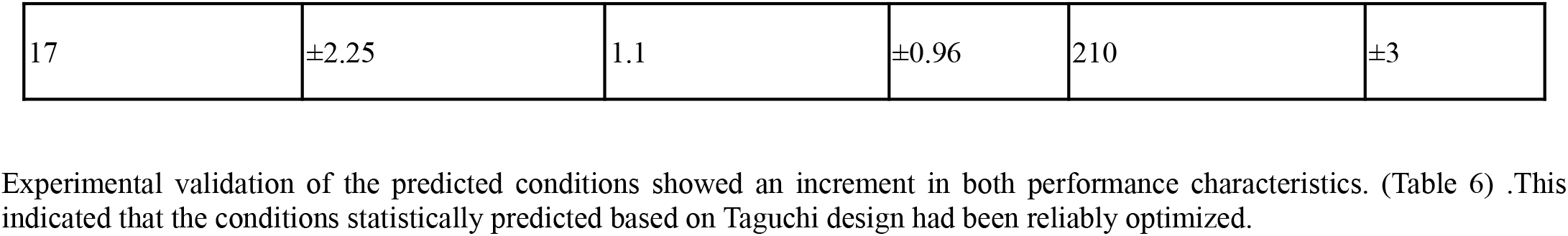
SD of different level of various factors for Yield (a) and EI% of BS (b)

| (a) |  |  |  |  |  |
|---|---|---|---|---|---|
| Carbon (%) | SD | Nitrogen (%) | SD | Agitation (rpm) | SD |
| 7.9 | ±0.05 | 0.16 | ±0.82 | 120 | ±1.87 |
| 14.78 | ±1.87 | 0.52 | ±1.50 | 180 | ±1.22 |
| 17 | ±0.60 | 1.1 | ±0.28 | 210 | ±1.64 |
| (b) |  |  |  |  |  |
| Carbon (%) | SD | Nitrogen (%) | SD | Agitation ( rpm ) | SD |
| 7.9 | ±1.41 | 0.16 | ±1.76 | 120 | ±6.51 |
| 14.78 | ±7.96 | 0.52 | ±4.48 | 180 | ±7.31 |
| 17 | ±2.25 | 1.1 | ±0.96 | 210 | ±3 |

Experimental validation of the predicted conditions showed an increment in both performance characteristics. (Table 6) .This indicated that the conditions statistically predicted based on Taguchi design had been reliably optimized.

**Table 6.** Increment in EI %, and Yield g/g wet biomass after experimental validation of predicted conditions for BS production.

|  | Predicted conditions | Experimentally validated conditions | Percent Increment |
|---|---|---|---|
| EI (%) of BS | 72.33 | 92.75 | 28.23% |
| Yield (g/g wet biomass) of BS | 1.13 | 1.28 | 13.27% |

**Table 7.** Activity of extract at 1X and 5X concentrations at different temperatures.

| **Temperature (C)** | **Emulsification Index** |  |
|---|---|---|
|  | **1X** | **5X** |
| -20 | 83% | 75% |
| 0 | 87.5% | 57.14% |
| 4-10 | 60% | 40% |
| 37 | 60% | 90% |
| 80 | 66.67% | VAPOURIZED |
| 121 | 100% | 100% |

The stability of BS produced showed tremendous capacity of endurance over a wide range of the parameters studied (Fig.5).Tolerance to extreme environmental conditions,of extracted BS from microbial sources has been reported by several workers in their studies (W. Jing et al., 2011; A.P.P. Santos et al., 2018; T. Udoh and J. Vinogradov, 2019).

**Fig. 5.**
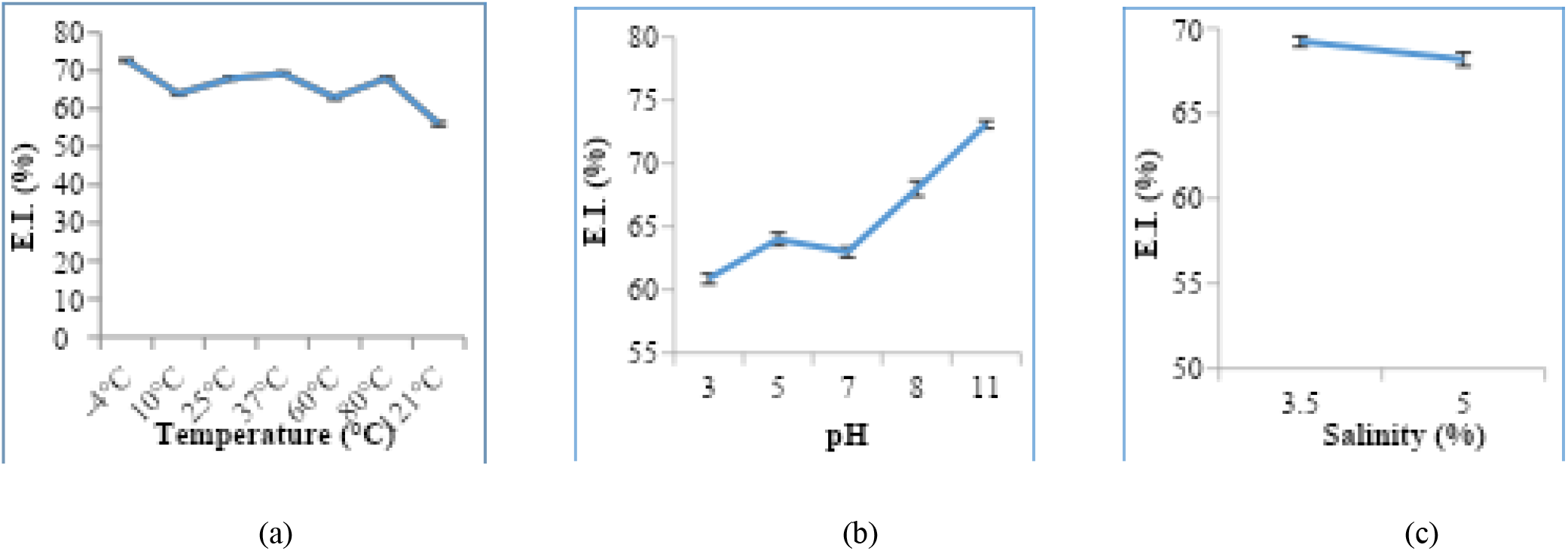
Effect of Temperature(a), pH (b) and salinity (c) on BS produced by *P. dendritiformis* in media amended with A.W.(at 10%)

**Figure 6.**
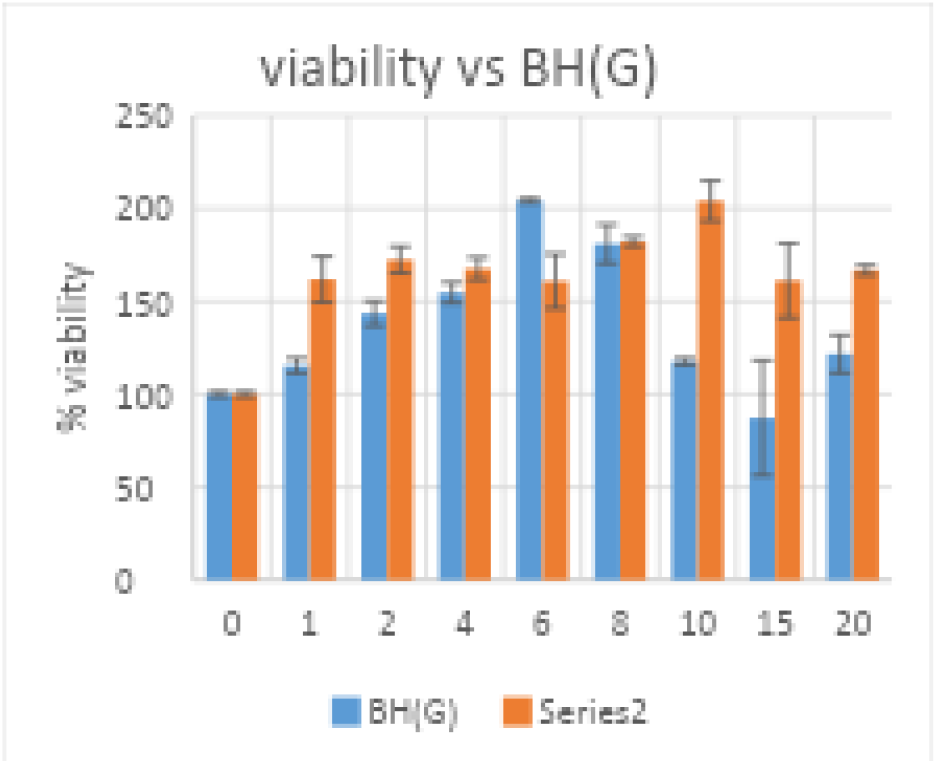
Effect of product on viability of cells after 1 hour and 48 hours respectively.

**Figure 7.**
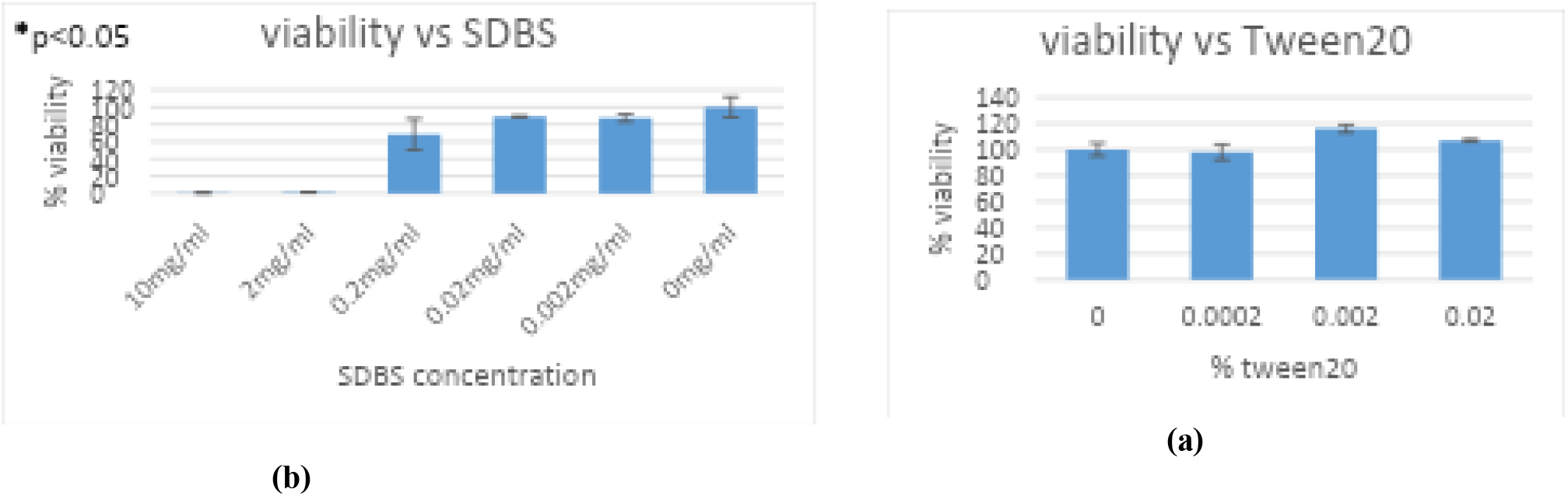
Viability of cells at various concentrations of SDBS(a) and Tween20(b)

### Stability of extracts

Greater activity was noted with 5X concentration of the extract with greater activity at higher temperature and alkaline pH values. At 1X concentration of extract, lower activity was observed but the activity was similarly better at higher temperatures and alkaline pH values.

### Toxicity test

The MTT assay indicated that the biosurfactant generated by *Paenibacillus dendritiformis* was substantially non-cytotoxic at the concentration range examined. Cell viability remained above the untreated control at concentrations of 1, 2, 4, 6, 8, 10 and 20, with viability ranging from 115.38% to 204.28%. Only at the concentration of 15 was there a decline below the control, with viability decreasing to 87.25%. Thus, the maximum reduction observed for the biosurfactant was only 12.75% and no concentration giving 50% loss in viability was observed in the tested range.

In contrast, conventional surfactants demonstrated significant decrease in cell viability with the increase in concentrations. At 0.2 mg/mL SDBS decreased the viability to 68.48% (31.52% reduction), whereas at 2 and 10 mg/mL, the viability was reduced to 1.20% (98.80% reduction) and 0.65% (99.35% reduction), respectively. Tween 20 exhibited a significant reduction at its highest tested concentration of 0.02% with a viability reduced to 46.18% (53.82% drop). The results obtained indicate that the biosurfactant has a comparatively lower cytotoxicity as compared to the evaluated chemical surfactants, confirming its potential as a lower-toxicity alternative.

Sodium Dodecylbenzene Sulfonate (SDBS) is a commonly used anionic surfactant having wetting, emulsifying, dispersing and detergent properties. It is microtoxic and has been recognised as a safe chemical raw material by the International Security Organisation. Tween 20 or Polysorbate 20 is a polysorbate-type nonionic surfactant. It’s stability and relative non-toxicity allows it to be used as a detergent and emulsifier in a number of domestic and pharmacological applications. Both are toxic at high concentrations.

## Conclusion

A 28.23% increase in performance characteristics of biosurfactant *viz*. EI of the biosurfactant and a 13.23 % increase of its yield of production was obtained when *Paenibacillus dendritiformis* was grown in an optimized medium containing carbon at 7.9 %, nitrogen at 1.1% provided by glucose 2%, ammonium nitrate 0.2%, mustard oil seed meal 10% in BH broth. The optimum agitation rate was found to be 180 r.p.m.

Experimental validation of predicted conditions conclusively established that the media components and conditions optimized in this study are reliable and can be used for commercial production of a highly stable biosurfactant.

## Acknowledgement

Financial assistance to Ms Gayatri Jeph as Rajiv Gandhi Fellowship (no.F.14-823(ST)/2010(SA-III) from the University Grants Commission, Government of India, is gratefully acknowledged. We acknowledge the University Supported Research Project Number GCU/RCC/2020-21/157/21 from The Maharaja Sayajirao University of Baroda and DBT-Boost to University Interdisciplinary Life Science Departments for Education and Research Programme (DBT-BUILDER)-Level III-Cat-III Programme (BT/INF/22/SP41403/2021) by the Department of Biotechnology for the ongoing work in Mayur Sahani’s lab (MSL). We acknowledge the support of the DST-PURSE Program (DST Reference No. SR/PURSE/2021/77), Department of Science and Technology, Government of India, for the establishment of the Sophisticated Instrumentation Facility (SIF) at Amity University Rajasthan, Jaipur.

